# Female variation in allocation of steroid hormones, antioxidants and fatty acids: a multilevel analysis in a wild passerine bird

**DOI:** 10.1101/467258

**Authors:** Lucia Mentesana, Caroline Isaksson, Wolfgang Goymann, Martin N. Andersson, Monika Trappschuh, Michaela Hau

## Abstract

The environment where an embryo develops can be influenced by components of maternal origin, which can shape offspring phenotypes and therefore maternal fitness. In birds that produce more than one egg per clutch, females differ in the concentration of components they allocate into the yolk along the laying sequence. However, identification of processes that shape female yolk allocation and thus offspring phenotype still remains a major challenge within evolutionary ecology. A way to increase our understanding is by acknowledging that allocation patterns can differ depending on the level of analysis, such as the population *versus* the among-female (within-population) level. We employed mixed models to analyze at both levels the variation in allocation along the laying sequence of four steroid hormones, three antioxidants, and four groups of fatty acids present in the egg yolks of wild great tits (*Parus major*). We also quantified repeatabilities for each component to study female consistency. At a population level, the concentrations/proportions of five yolk components varied along the laying sequence, implying that the developmental environment is different for offspring developing in first *versus* last eggs. Females varied substantially in the mean allocation of components and in their plasticity along the laying sequence. For most components, these two parameters were negatively correlated. Females were also remarkably repeatable in their allocation. Overall, our data emphasize the need to account for female variation in yolk allocation along the laying sequence at multiple levels, as variation at a population level is underpinned by different individual patterns. Our findings also highlight the importance of considering both levels of analysis in future studies investigating the causes and fitness consequences of yolk compounds. Finally, our results on female repeatability confirm that analyzing one egg per nest is a suitable way to address the consequences of yolk resource deposition for the offspring.

## Introduction

Female birds can influence the physiological conditions in which their embryos will develop by differential allocation of resources into the egg yolk, thus generating variation in offspring phenotype and influencing fitness (Mousseau and Fox 1998). Such resources can be hormones (Schwabl 1993) and nutrients (Surai 2002, Hulbert and Abbott 2011), which can affect embryonic growth and development and also provide protection to oxidative damage. In bird species that produce more than one egg per clutch, females often vary the concentration of components they allocate along the laying sequence (e.g., Royle et al. 1999, Blount et al. 2002, Hõrak et al. 2002, Saino et al. 2002, Tschirren et al. 2004, Bourgault et al. 2007, Rubolini et al. 2011, Lessels et al. 2016, Toledo et al. 2016). Hence, depending on the egg in which they develop, offspring from the same clutch can be exposed to different environments during embryonic development. Despite maternal effects being an important factor in evolution (Mousseau and Fox 1998), the identification of the processes that shape female yolk allocation still remains a major challenge in biology.

A way to increase our understanding of yolk allocation is by acknowledging that females can show different patterns of allocation along the laying sequence when analyzed at multiple levels (Meyers and Bull 2002). This phenotypic plasticity (defined as the property of a given genotype to produce different phenotypes as environmental conditions change; Via et al. 1995, Pigliucci 2001, Nussey et al. 2007) observed along the laying sequence can be analyzed at a population level by looking at the average female allocation of components along the laying sequence (Figure 1a). At a population level, consistent patterns of average yolk allocation along the laying sequence have been reported for several bird species. For example, domestic canaries (*Serinus canaria*) and black-headed gulls (*Chroicocephalus ridibundus*) increase the concentration of androgens, while zebra finches (*Taeniopygia guttata*) decrease the concentration of these components from the first to the last egg (reviewed by Groothuis et al. 2005, Gil 2008). This population-level variation, hereafter referred to as “mean phenotypic plasticity”, has been interpreted in light of sibling competition and parent-offspring conflict (Müller et al. 2007). However, a lack of consistency across populations of the same species has also been reported, for example in great tits (*Parus major;* Tschirren et al. 2004, Groothuis et al. 2008, Lessels et al. 2016). This inconsistency indicates that the patterns of allocation are more complex than previously assumed.

**Figure 1.**
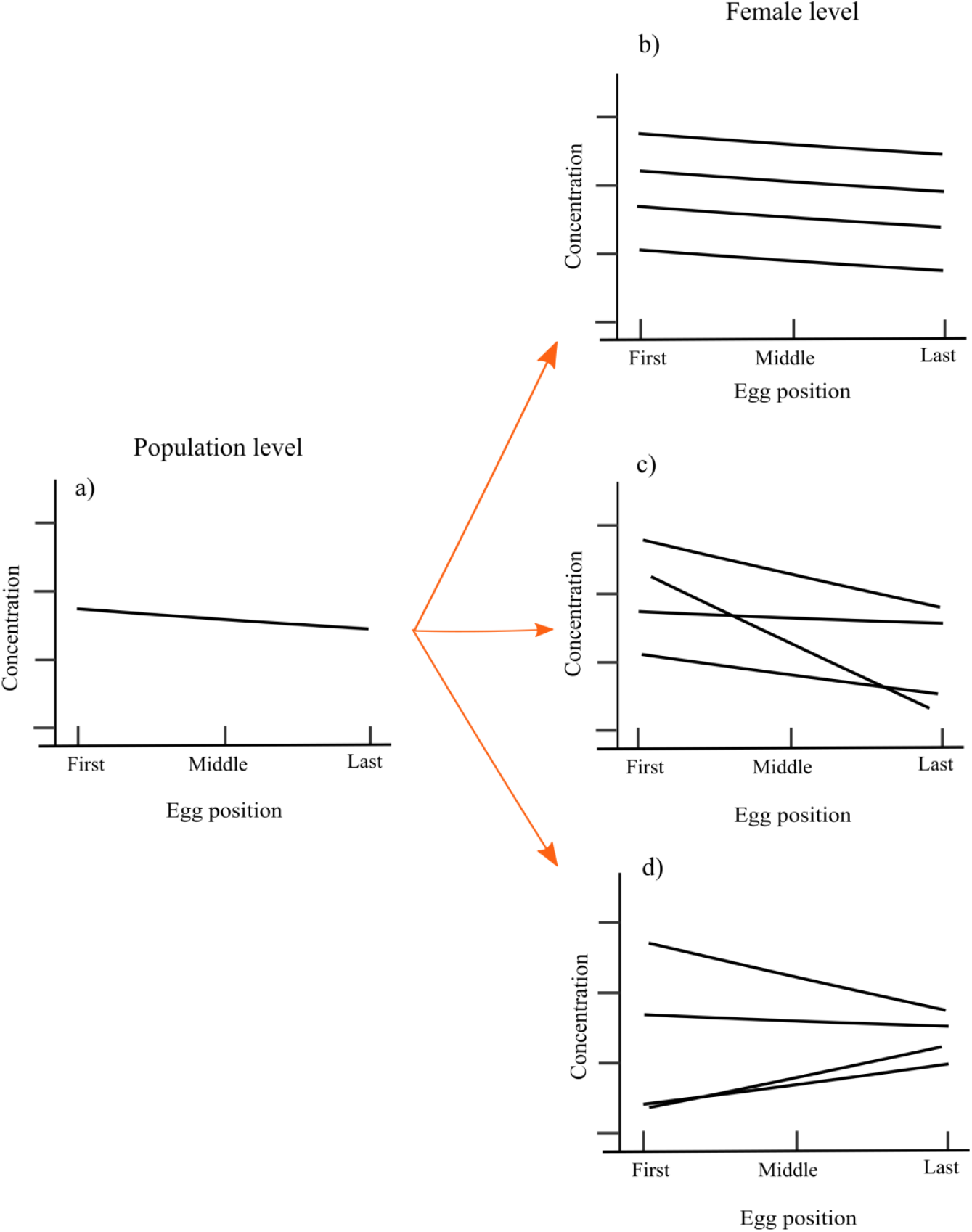
Schematical depiction of the two levels of analysis used in this study to understand female yolk allocation in avian species that produce clutches of more than one egg. (a) The mean phenotypic plasticity (i.e., average population variation) along the laying sequence can be caused by variation among females. (b) Females (individuals represented by different solid lines) can vary in the mean allocation of yolk components, (c) in the individual phenotypic plasticity (i.e., slope of allocation along the laying sequence), or (d) in both the mean and slope of allocation. In the latter case, the two parameters can also be positively or negatively correlated. For example, females that on average allocate a high concentration of a specific yolk component will more strongly decrease the concentration of that component along the laying sequence ((d); upper line).

Phenotypic plasticity in allocation patterns at a population level might not necessarily provide information regarding individual-level variation (Nussey et al. 2007, Dingemanse et al. 2010). The same pattern observed at a population level might be driven by females differing in the mean allocation of components (Figure 1b), in the slope of changes along the laying sequence (hereafter referred to as “individual phenotypic plasticity”; Figure 1c), or in both parameters which covary (Figure 1d). Mixed results across populations of the same species may therefore result from differences among females in allocation patterns within each population (Nussey et al. 2007, Dingemanse et al. 2010). Individual female variation can be quantified by using a reaction norm approach, which allows us to estimate how much of each yolk component females transfer on average into the yolk (i.e., the elevation of the reaction norm), the change in female allocation along the laying sequence (i.e., individual phenotypic plasticity), as well as the covariation between elevation and slope (Nussey et al. 2007, Dingemanse et al. 2010, Dingemanse and Wolf 2013). In particular, the presence of correlations between the mean allocation of components and the individual phenotypic plasticity suggests that maternal effects cannot be fully evaluated by studying each of these components separately, since selection could be acting on each of these sources of variation and/or directly on their correlation. Since natural selection operates at the individual level, accounting for individual female variation (i.e., mean allocation, phenotypic plasticity and the correlation between these two parameters) in yolk allocation along the laying sequence is therefore of key importance if we aim to understand the evolutionary causes and consequences of variation in female allocation.

Here we employed linear mixed models to study the allocation along the laying sequence at both population- and among female-levels in the yolks of 11 clutches of freshly laid eggs collected from a wild population of great tits. Our emphasis in this investigation was to provide new insights into individual-level patterns. Furthermore, since the majority of research to date has focused on androgens, here we also aim at simultaneously quantifying additional yolk components to increase our understanding of the factors contributing to such differences in allocation patterns observed along the laying sequence. For each egg collected we measured the concentrations of four steroid hormones (the androgens androstenedione, 5α-dihydrotestosterone, testosterone, and the glucocorticoid corticosterone), three antioxidants (vitamin E, lutein, zeaxanthin) and the proportions of four groups of fatty acids (saturated fatty acids, monounsaturated fatty acids, omega (ω)-3 polyunsaturated fatty acids (PUFA), ω-6 PUFA). We then quantified female consistency in allocation for each yolk component (e.g., whether females that allocate on average high levels of a yolk component always allocate high concentrations along the laying sequence compared to other females) by calculating adjusted repeatabilities.

We selected specific yolk components for our analysis because of possible interactive effects on offspring phenotype, although the statistical determination of such possible interactions is beyond the scope of the current study. Steroid hormones such as androgens and glucocorticoids can enhance offspring growth, competitive ability and survival, while also possibly causing immunosuppression and oxidative stress (Schwabl 1993, Groothuis and Schwabl 2002, Groothuis et al. 2005, Gil 2008, Groothuis and Schwabl 2008, Haussmann et al. 2012, Treidel et al. 2013). Conversely, antioxidants like the carotenoids and vitamin E can enhance the immune system and mitigate oxidative stress caused by embryo growth (Surai 2000, Saino et al. 2003, Yigit et al. 2014, Parolini et al. 2017, Watson et al. 2018). Fatty acids, in turn, provide the avian embryo with almost all the energy and building blocks required to sustain development within the egg (Noble and Cocchi 1990, Surai and Speake 2008). In particular, PUFAs are vital components for the formation of cell membranes, heart functioning and brain development (Hulbert and Abbott 2011). However, highly unsaturated PUFAs are susceptible to lipid peroxidation through reactive oxygen species that are generated by embryonic metabolism (Pamplona et al. 2002, Larsson et al. 2004, Hulbert and Abbott 2011, Yigit et al. 2014). Furthermore, we recorded lay date, ambient temperatures at the time of laying, and female body condition. Factors such as lay date and ambient temperature may explain the concentration/proportion of some yolk components, especially those known to be highly influenced by the quality and quantity of food consumed by the mother (antioxidants like vitamin E and carotenoids and PUFAs; Surai and Speake 2008; Hulbert and Abbott 2011). Internal variables such as female body condition are also usually considered important factors for determining the yolk allocation of substances like hormones (reviewed by Groothuis et al. 2005), antioxidants (Blount et al. 2002, Williamson et al. 2006) and fatty acids (Raclot 2003, Price et al. 2008).

## Methods

### Study species, field site and sampling

Great tits are small passerine birds that breed inside cavities. Females usually lay one egg each day early in the morning. While having a seed-dominated winter diet, great tits feed primarily on caterpillars during the breeding season (Royama 1970) which are an important source of proteins and fatty acids (Isaksson and Andersson 2007, Andersson et al. 2015, Isaksson et al. 2015).

For this study, fresh laid eggs of entire clutches were collected in April and May 2015 from a nest box population of great tits in the Dellinger Buchet, a mosaic of deciduous and coniferous forest in Southern Germany (Bavaria; 48°03’ N, 11°13’ E, 620 m above sea level). The breeding stage was monitored every second day from the first signs of nest construction onwards. Eggs were collected on the date of lay between 8:00 and 13:00h and each removed egg was replaced with a dummy egg. In total, 93 eggs from 11 first clutches were collected. Clutch sizes ranged from 6 to 10 eggs (mean ± SD: 8.45 ± 1.13 eggs). Once in the laboratory, egg measurements were taken following established protocols by Lessells et al. (2002). Briefly, eggs were weighed, opened, and the yolk separated from the remainder of the egg. Excess albumen was removed from the yolk by rolling it on a piece of paper. During this step the vitelline membrane of the yolk got disrupted in three out of 93 eggs, but since the amount of yolk lost was minor, we included these eggs in the analysis. The yolk was weighed, homogenized in distilled water and immediately stored at −80°C until further analysis. The egg shell was washed, weighed and dried at room temperature until the next morning when the dry shell was weighed again. Albumen mass was estimated by subtracting the wet yolk mass and the dry shell mass from the total egg weight.

We measured local environmental conditions during the formation of the eggs. Ambient temperature was recorded every hour via i-buttons (DS9093A+ Thermochron iButton) placed in 12 different places across the study site. For our analyses, we used the mean temperature on the three days preceding the lay date of each egg, hereafter referred as mean temperature, since the phase of follicular growth lasts about three days in great tits (Walsberg 1983). Preliminary exploration of our data showed that other environmental factors like rainfall did not have an effect on the allocation of yolk components (results not shown). Hence, we only included mean ambient temperature in our analyses.

Nine of the 11 females were captured when incubating the replaced eggs, on average 4.78 ± 1.62 days after they had laid their last egg. Females were marked with a numbered aluminum ring and plastic split rings with a unique colour combination for individual identification. All females were adults (> 1 year), as determined from plumage characteristics. Body mass (to the nearest 0.1 g), tarsus length (to the nearest 0.1 mm) and wing length (to the nearest 0.1 mm) were measured for each captured individual. Scaled body mass index was used as an indicator of female condition, since it accounts for the allometric relationship between different measures of body size and mass (Peig and Green 2009). As recommended by Peig and Green (2009), parameters with the highest correlation in our population were used to estimate female condition: body mass and wing length (*r* = 0.68; p-value = 0.05).

### Steroid hormone analysis

To quantify androstenedione, 5α-dihydrotestosterone, testosterone, and corticosterone concentrations, we conducted radioimmunoassays following the method described by Wingfield and Farner (1976), modified by Goymann et al. (2008) with additional adjustments for the measurement of egg yolk following Schwabl (1993). Steroids were extracted from the yolk in two sets of assays. Because one of the key aspects of our manuscript was to study yolk allocation of individual females, and variation in yolk allocation among females is typically higher than within females (reviewed by Groothuis et al. 2005), eggs from the same clutch were always included in the same assay. On average, 50 µl of the yolk/water emulsion was transferred to 16 x 100 glass test tubes. Along with the samples, two blanks containing 300 µl distilled water and three positive controls containing 100 µl stripped chicken plasma pools were also prepared. Distilled water was added to all tubes to have the same final volume (300 µl). Next, we added 10 µl tritiated steroid (1500 dpm; PerkinElmer, MA, USA) of all steroid hormones to be measured to all tubes except the blanks to estimate extraction efficiency. We then added 4 ml of diethyl ether to each sample. After overnight equilibration the samples were centrifuged, the supernatant was collected and dried under a nitrogen stream in a water bath at 40 °C. Each sample was then subjected to a second extraction by adding 2 ml dichloromethane. The dried supernatant was re-suspended in 1 ml 99% ethanol. After an overnight reconstitution, extracts were centrifuged, the supernatant collected and again dried under a nitrogen stream in a water bath at 40 °C, and reconstituted in isooctane with 2% ethyl acetate. Steroids were then further separated via diatomaceous earth column chromatography. Fractions containing different steroids were eluted by mixing ethyl acetate (EtAc) with isooctane at increasing concentrations (2%, 10%, 25%, and 45% EtAc for androstenedione, 5α-dihydrotestosterone, testosterone and corticosterone, respectively). Each eluted fraction was collected in 12 × 75 mm glass tubes and evaporated under a nitrogen stream. Androgens (androstenedione, 5α-dihydrotestosterone, and testosterone) were re-dissolved in 300 µl phosphate-buffered saline. 80 µl of the resuspended fraction were used to estimate individual extraction recoveries. Recoveries for the two sets of columns (n = 50 samples per set) were within the expected range previously reported for great tits (Tschirren et al. 2004, Groothuis et al. 2008, Lessels et al. 2016) and mean ± SD were as follows: androstenedione = 83 ± 2.83%, 5α-dihydrotestosterone = 55% (in both sets), testosterone = 51 ± 1.41%. Duplicates of 100 µl were used for the radioimmunoassays. The hormone concentration of each sample was corrected for the individual extraction efficiency. Some of the samples had concentrations above the upper detection limit of the assay (which was at 200 pg per tube), hence we extracted another proportion of the yolk following the same protocol but using a smaller volume (70 µl) of the extracted sample for the radioimmunoassays (androstenedione: 79 samples; testosterone: 6 samples). Recoveries for these two additional sets of columns were comparable to the first set of assays (mean ± SD only for androstenedione): androstenedione = 78.5 ± 0.71%, testosterone = 89%. Polyclonal antibodies used were the following: AN6-22 for androstenedione, DT3-351 for 5α-dihydrotestosterone and T3-125 for testosterone (all Esoterix Endocrinology, Inc., CA, USA). The lower detection limit was 0.80 pg/ml for androstenedione, 0.56 pg/ml for 5α-dihydrotestosterone, and 0.36 pg/ml for testosterone. Blanks were all below detection limits. All samples were analyzed in two assays. The intra-assay variations for both assays, determined from the three positive controls, were: androstenedione = 7.65 ± 0.85%, 5α-dihydrotestosterone = 4.15 ± 0.25%, testosterone = 9.3 ± 0.5%. The inter-assay variations, determined by including the first positive control per assay, were (mean ± SD): androstenedione = 16.2%, 5α-dihydrotestosterone = 3.4%, and testosterone = 12.4%. For those samples that needed to be re-extracted the intra-assay variations were: androstenedione (mean ± SD) = 8.65 ± 1.13%, testosterone = 4.1%.

Corticosterone concentrations were determined by enzyme immunoassay (Lot No: 12041402D and 08241511, Enzo Life Sciences GmbH, Germany). After column chromatography, the corticosterone fractions of the samples were dried and then dissolved in 350 µl assay buffer. An aliquot of 80 µl was used to estimate individual extraction recoveries (mean ± SD recoveries were 57 ± 11.31 %). Duplicates of 100 µl were added to individual wells and samples were distributed across 4 assays. The intra-assay coefficients of variation were 1.9%, 5.5%, 2.4%, 2.3% (calculated from two replicate standards per plate) and the inter-assay variation was 3%. The corticosterone antibody has a 0.5% cross-reactivity with progesterone, but the column chromatographic separation of steroid hormones had separated corticosterone from progesterone (Wingfield and Farner 1976), so that cross-reactivity can be excluded.

### Antioxidant extraction and HPLC analysis

Vitamin E (α-tocopherol) and carotenoids (lutein and zeaxanthin) were extracted simultaneously. Briefly, to 20 mg yolk, 200 µl acetone with internal standard - 600 µM retinyl acetate and 1 mM tocopheryl acetate (Sigma-Aldrich, Stockholm, Sweden) was added, followed by vortexing. The samples were then left overnight at −80°C. The following day, 200 µl tert-butyl methyl ether was added, followed by vortexing. Samples were centrifuged at 10°C for 5 min (13,000 rpm), and the supernatant was transferred to a new tube and dried under nitrogen gas. The samples were washed twice with 200 µl acetone, followed by vortexing and centrifugation; the supernatant was removed to a new tube between the washes. Again, samples were dried under nitrogen gas. The residue was dissolved in 100 µl methanol-acetonitrile (30:70). The amount of α-tocopherol, lutein and zeaxanthin were determined by high performance liquid chromatography (HPLC) with the following specifications: column Phenomenex Syndergi 4u Hydro-RP 80A, 250 × 3 mm + 4 × 2 mm guard column, isocratic 20 % MeOH, 80 % AcCN, 12 min, 1,2 ml/min, oven 40 °C, injection 5 µl, UV 450 nm, FL ex: 290 nm, em: 325 nm. The concentrations were calculated from standard curves made from lutein, zeaxanthin and tocopherol, along with corrections for their respective internal standards.

### Fatty acid extraction and quantification

The fatty acids were extracted as described by Eikenaar et al. (2017). Briefly, a total lipid extraction of approximately 5 mg of yolk was done using chloroform and methanol (2:1 v/v). Base methanolysis was carried out to transform the fatty acids into corresponding fatty acid methyl esters (FAMEs). The FAMEs were extracted using heptane (>99%; VWR Prolabo), and the extracts were analyzed using an Agilent 5975 mass spectrometer coupled to an Agilent 6890 gas chromatograph with an HP-INNOWax PEG column (30 m, 0.25 mm i.d., 0.25 mm film thickness; Agilent). Analyses and quantification of chromatograms were performed using ChemStation software (Agilent). FAMEs were identified by comparing mass spectra and retention times with those of synthetic standards (Supelco 37-Component FAME Mix, Sigma-Aldrich).

### Data handling and statistical analysis

In total, we studied 11 yolk components. For yolk steroid hormones and antioxidants, the statistical analyses were done using the concentration (in pg/ml or standard micromolar, respectively), while for fatty acids we analyzed the proportion of each fatty acid group (Andersson et al. 2015, Isaksson et al. 2017). For simplicity, lutein and zeaxanthin were pooled and referred to as total carotenoids whenever both components showed similar results. Fatty acid proportions were calculated by dividing the peak area of each fatty acid by the sum of the peak areas of all fatty acids in each individual sample (Andersson et al. 2015, Isaksson et al. 2017). The proportions of all individual fatty acids within a certain chemical class of fatty acid were then combined to obtain relative levels of total saturated fatty acids, total monounsaturated fatty acids, total ω-3 PUFA, and total ω-6 PUFA (Andersson et al. 2015, Isaksson et al. 2017). Furthermore, one of our aims was to study the difference in yolk components along the laying sequence. Since great tits vary in clutch size, analyzing yolk components in terms of only egg number is inadequate. We therefore used the relative egg position (determined as egg position/N eggs per clutch) within a range from 0 to 1 as a standardized variable in our analysis (indicated as ‘first’, ‘middle’ and ‘last’ in the figures for illustrative purposes only).

We ran three univariate mixed-models fitting each yolk component or group of yolk component, respectively, as a response variable. All continuous explanatory variables were mean-centered and their variance standardized to facilitate comparison of variance components across traits. First, to study the relationship between yolk components and female body condition we used the scaled body mass index as a measure of female condition, egg position, and mean ambient temperature (temperature range = 7.6 – 16.8°C) as covariates. Lay date was not included in this analysis to avoid overparametrization. Despite the fact that we were interested in the effect that female body condition has on yolk components (i.e., fixed factor), female identity and the interaction between female identity and laying sequence were fitted in the model as random elevation and random slope, respectively (for a further discussion of the rationale of this approach see Schielzeth and Forstmeier, 2009).

Second, to study female allocation at the population and individual level, we used a random regression model (i.e., a reaction norm framework; Nussey et al. 2007, Dingemanse et al. 2010). Egg position, lay date (date of first egg, range = 16^th^ of April – 9^th^ of May) and mean ambient temperature were fitted as covariates (the correlation between laying date and mean ambient temperature was relatively low; *r* = 0.26; p-value = 0.01). Female identity (i.e., random elevation) and the interaction between female identity with respect to laying (i.e., random slope) were also fitted in the model. By using this random elevation-slope approach we were able to estimate three parameters: i) the among-female variation in the average concentration/proportion of egg components (i.e., variance in elevation; Figure 1b), ii) the among-female variation in plasticity along the laying sequence (i.e., variance in slope; Figure 1c), and iii) the correlation between elevation and slope (Figure 1d).

Finally, to calculate the repeatability (*R*) of yolk components among females, we built a model similar to the one described in the previous paragraph, but only including female identity as a random intercept. The repeatability of female yolk allocation was calculated as the variance component explained by female identity divided by the total variance (female identity + residual) in the presence of fixed effects (“adjusted repeatability”; Nakagawa and Schielzeth 2010; i.e., egg position, lay date and mean ambient temperature).

All statistical analyses were performed in R statistical freeware R-3.3.3 (R Core Team 2017) using the “lme4” and “arm” packages in a Bayesian framework with non-informative priors. We assumed a Gaussian error distribution, which was confirmed for all response variables after visual inspection of model residuals. When necessary, response variables were transformed (details on transformations are provided in the Tables). We subsequently used the *sim* function to simulate values from the posterior distributions of model parameters. We extracted the 95% Bayesian Credible Interval (CrI) around the mean (Gelman and Hill 2007), and assessed statistical support by obtaining the posterior distribution of each parameter. CrI provide more valuable information than p-values, like for example, the uncertainty around the estimates. We use the term “meaningful effect” if zero was not included within the 95% CrI (Korner-Nievergelt et al. 2015). For intervals overlapping zero only slightly, we report the posterior probability of the estimate being positive or negative (see Korner-Nievergelt et al. 2015 for further discussion of how to infer conclusions from Bayesian statistical analysis).

## Results

### Egg yolk components, environmental effects and female body condition

Egg mass increased along the laying sequence by 12%, while yolk mass only tended to increase (Supplementary material Appendix 1, Table A1). The concentrations of androstenedione, 5α-dihydrotestosterone and testosterone were substantially higher than those of corticosterone (Supplementary material Appendix 1, Table A2). Among the antioxidants, the carotenoid lutein was the most abundant, followed by zeaxanthin and vitamin E. Furthermore, 20 fatty acids were identified (Supplementary material Appendix 1, Table A3), with the monounsaturated fatty acid group contributing most to the total fatty acid content (around 46%; Supplementary material Appendix 1, Table A2), followed by saturated fatty acids, ω-6 PUFAs and lastly ω-3 PUFAs.

Females that laid eggs later in the breeding season allocated higher concentrations of androstenedione and higher proportions of saturated fatty acids and ω-3 PUFAs into the yolk, while the concentration of corticosterone and the proportion of ω-6 PUFAs decreased over the breeding season (Table 1). The mean ambient temperature had a positive effect on the proportion of ω-6 PUFAs, and a negative one on the concentrations of corticosterone and the proportion of monounsaturated fatty acids (Table 1). Neither lay date nor mean ambient temperature influenced the concentrations of yolk antioxidants. Female scaled body mass index had a negative effect on 5α-dihydrotestosterone concentrations and a positive one on vitamin E concentrations in the yolk (Supplementary material Appendix 1, Table A4). However, scaled body mass index had a weak effect, if any, on most of the other yolk components.

**Table 1.**
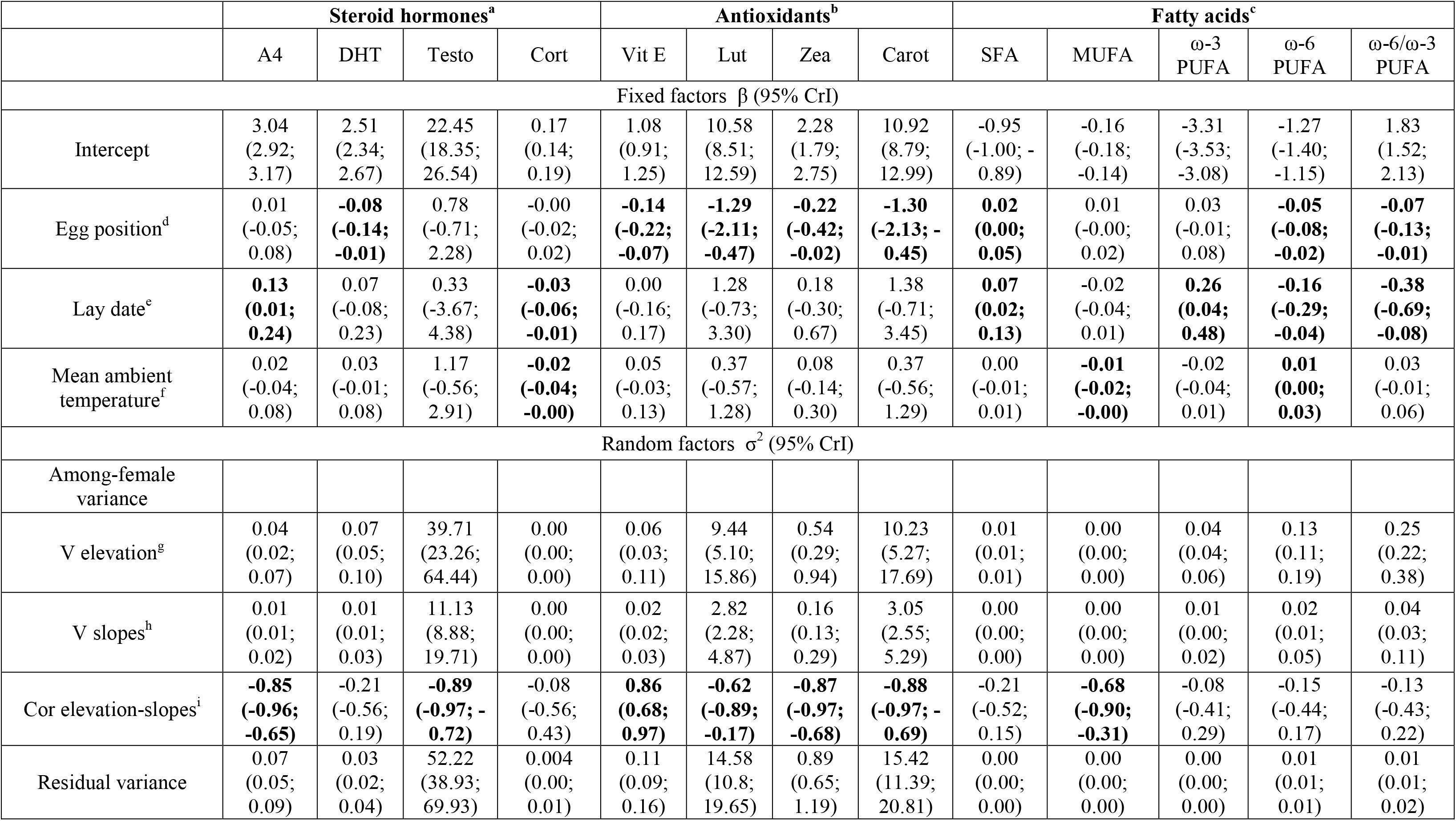
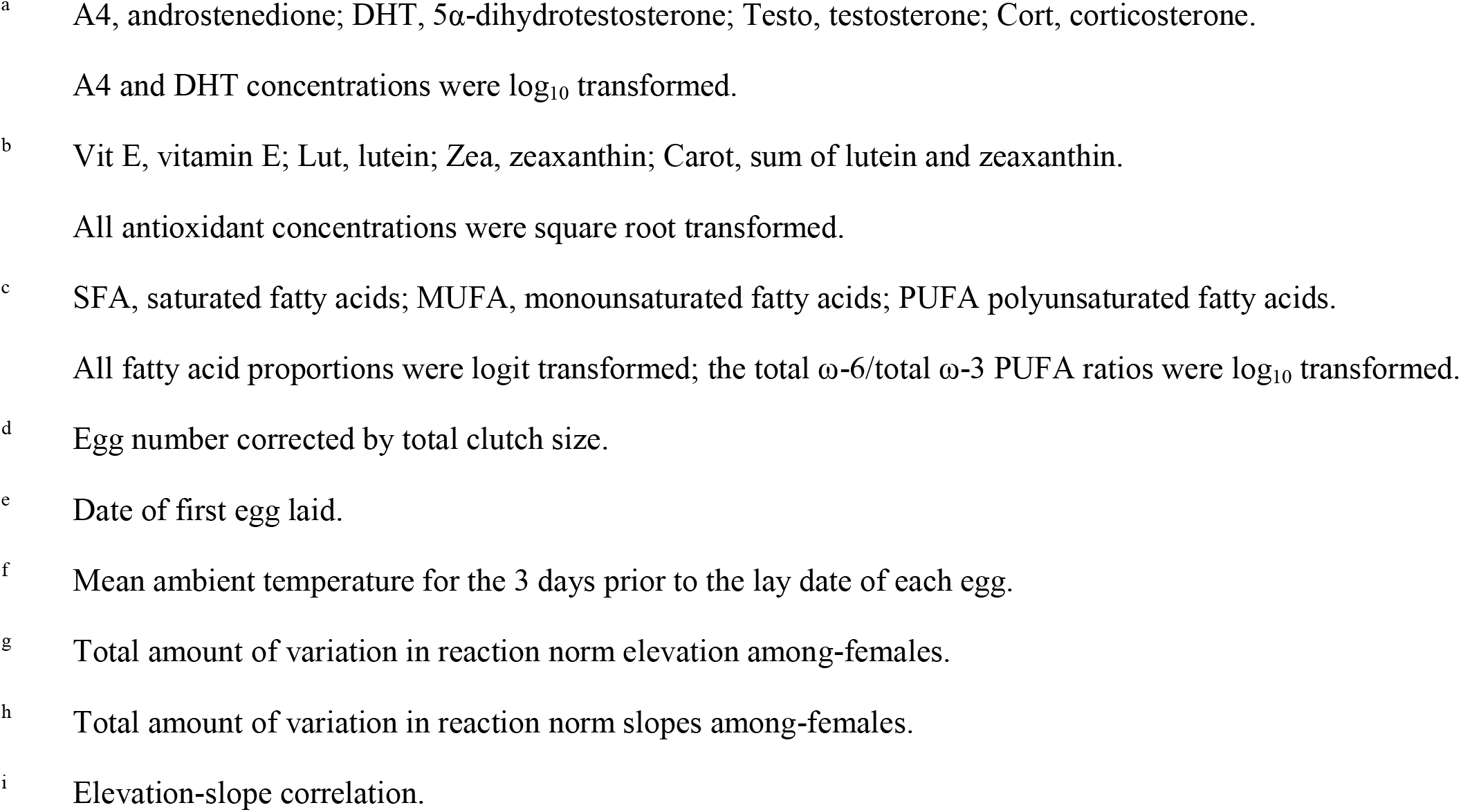
Results from linear mixed-effects models estimating fixed and random effects to explain variation in yolk components and variation among females. Egg position, lay date and mean ambient temperature were fitted as covariates, and random slopes were fitted for female identity with respect to egg position. We present fixed (β) and random (σ^2^) parameters with their 95% credible intervals (CrI) in brackets. All explanatory variables were mean centered; hence the intercepts refer to the average value of covariates. Fixed factors with a statistically meaningful effect (i.e., if zero is not included within the 95% CrI) are presented in bold. Estimates and CrI of ‘0.00’ represent an effect smaller than 0.01. ^a^ A4, androstenedione; DHT, 5á-dihydrotestosterone; Testo, testosterone; Cort, corticosterone.A4 and DHT concentrations were log10 transformed. ^b^ Vit E, vitamin E; Lut, lutein; Zea, zeaxanthin; Carot, sum of lutein and zeaxanthin. All antioxidant concentrations were square root transformed. ^c^ SFA, saturated fatty acids; MUFA, monounsaturated fatty acids; PUFA polyunsaturated fatty acids. All fatty acid proportions were logit transformed; the total ù-6/total ù-3 PUFA ratios were log10 transformed. ^d^ Egg number corrected by total clutch size. ^e^ Date of first egg laid. ^f^ Mean ambient temperature for the 3 days prior to the lay date of each egg. ^g^ Total amount of variation in reaction norm elevation among-females. ^h^ Total amount of variation in reaction norm slopes among-females. ^i^ Elevation-slope correlation.

|  | Steroid hormones<sup>a</sup> |  |  |  | Antioxidants<sup>b</sup> |  |  |  | Fatty acids<sup>c</sup> |  |  |  |  |
| --- | --- | --- | --- | --- | --- | --- | --- | --- | --- | --- | --- | --- | --- |
|  | A4 | DHT | Testo | Cort | Vit E | Lut | Zea | Carot | SFA | MUFA | ω-3<br>PUFA | ω-6<br>PUFA | ω-6/ω-3<br>PUFA |
| Fixed factors β (95% CrI) |  |  |  |  |  |  |  |  |  |  |  |  |  |
| Intercept | 3.04<br>(2.92;<br>3.17) | 2.51<br>(2.34;<br>2.67) | 22.45<br>(18.35;<br>26.54) | 0.17<br>(0.14;<br>0.19) | 1.08<br>(0.91;<br>1.25) | 10.58<br>(8.51;<br>12.59) | 2.28<br>(1.79;<br>2.75) | 10.92<br>(8.79;<br>12.99) | -0.95<br>(-1.00; -<br>0.89) | -0.16<br>(-0.18;<br>-0.14) | -3.31<br>(-3.53;<br>-3.08) | -1.27<br>(-1.40;<br>-1.15) | 1.83<br>(1.52;<br>2.13) |
| Egg position<sup>d</sup> | 0.01<br>(-0.05;<br>0.08) | **-0.08**<br>**(-0.14;**<br>**-0.01)** | 0.78<br>(-0.71;<br>2.28) | -0.00<br>(-0.02;<br>0.02) | **-0.14**<br>**(-0.22;**<br>**-0.07)** | **-1.29**<br>**(-2.11;**<br>**-0.47)** | **-0.22**<br>**(-0.42;**<br>**-0.02)** | **-1.30**<br>**(-2.13; -**<br>**0.45)** | **0.02**<br>**(0.00;**<br>**0.05)** | 0.01<br>(-0.00;<br>0.02) | 0.03<br>(-0.01;<br>0.08) | **-0.05**<br>**(-0.08;**<br>**-0.02)** | **-0.07**<br>**(-0.13;**<br>**-0.01)** |
| Lay date<sup>e</sup> | **0.13**<br>**(0.01;**<br>**0.24)** | 0.07<br>(-0.08;<br>0.23) | 0.33<br>(-3.67;<br>4.38) | **-0.03**<br>**(-0.06;**<br>**-0.01)** | 0.00<br>(-0.16;<br>0.17) | 1.28<br>(-0.73;<br>3.30) | 0.18<br>(-0.30;<br>0.67) | 1.38<br>(-0.71;<br>3.45) | **0.07**<br>**(0.02;**<br>**0.13)** | -0.02<br>(-0.04;<br>0.01) | **0.26**<br>**(0.04;**<br>**0.48)** | **-0.16**<br>**(-0.29;**<br>**-0.04)** | **-0.38**<br>**(-0.69;**<br>**-0.08)** |
| Mean ambient<br>temperature<sup>f</sup> | 0.02<br>(-0.04;<br>0.08) | 0.03<br>(-0.01;<br>0.08) | 1.17<br>(-0.56;<br>2.91) | **-0.02**<br>**(-0.04;**<br>**-0.00)** | 0.05<br>(-0.03;<br>0.13) | 0.37<br>(-0.57;<br>1.28) | 0.08<br>(-0.14;<br>0.30) | 0.37<br>(-0.56;<br>1.29) | 0.00<br>(-0.01;<br>0.01) | **-0.01**<br>**(-0.02;**<br>**-0.00)** | -0.02<br>(-0.04;<br>0.01) | **0.01**<br>**(0.00;**<br>**0.03)** | 0.03<br>(-0.01;<br>0.06) |
| Random factors σ<sup>2</sup> (95% CrI) |  |  |  |  |  |  |  |  |  |  |  |  |  |
| Among-female<br>variance |  |  |  |  |  |  |  |  |  |  |  |  |  |
| V elevation<sup>g</sup> | 0.04<br>(0.02;<br>0.07) | 0.07<br>(0.05;<br>0.10) | 39.71<br>(23.26;<br>64.44) | 0.00<br>(0.00;<br>0.00) | 0.06<br>(0.03;<br>0.11) | 9.44<br>(5.10;<br>15.86) | 0.54<br>(0.29;<br>0.94) | 10.23<br>(5.27;<br>17.69) | 0.01<br>(0.01;<br>0.01) | 0.00<br>(0.00;<br>0.00) | 0.04<br>(0.04;<br>0.06) | 0.13<br>(0.11;<br>0.19) | 0.25<br>(0.22;<br>0.38) |
| V slopes<sup>h</sup> | 0.01<br>(0.01;<br>0.02) | 0.01<br>(0.01;<br>0.03) | 11.13<br>(8.88;<br>19.71) | 0.00<br>(0.00;<br>0.00) | 0.02<br>(0.02;<br>0.03) | 2.82<br>(2.28;<br>4.87) | 0.16<br>(0.13;<br>0.29) | 3.05<br>(2.55;<br>5.29) | 0.00<br>(0.00;<br>0.00) | 0.00<br>(0.00;<br>0.00) | 0.01<br>(0.00;<br>0.02) | 0.02<br>(0.01;<br>0.05) | 0.04<br>(0.03;<br>0.11) |
| Cor elevation-slopes<sup>i</sup> | **-0.85**<br>**(-0.96;**<br>**-0.65)** | -0.21<br>(-0.56;<br>0.19) | **-0.89**<br>**(-0.97; -**<br>**0.72)** | -0.08<br>(-0.56;<br>0.43) | **0.86**<br>**(0.68;**<br>**0.97)** | **-0.62**<br>**(-0.89;**<br>**-0.17)** | **-0.87**<br>**(-0.97;**<br>**-0.68)** | **-0.88**<br>**(-0.97; -**<br>**0.69)** | -0.21<br>(-0.52;<br>0.15) | **-0.68**<br>**(-0.90;**<br>**-0.31)** | -0.08<br>(-0.41;<br>0.29) | -0.15<br>(-0.44;<br>0.17) | -0.13<br>(-0.43;<br>0.22) |
| Residual variance | 0.07<br>(0.05;<br>0.09) | 0.03<br>(0.02;<br>0.04) | 52.22<br>(38.93;<br>69.93) | 0.004<br>(0.00;<br>0.01) | 0.11<br>(0.09;<br>0.16) | 14.58<br>(10.8;<br>19.65) | 0.89<br>(0.65;<br>1.19) | 15.42<br>(11.39;<br>20.81) | 0.00<br>(0.00;<br>0.00) | 0.00<br>(0.00;<br>0.00) | 0.00<br>(0.00;<br>0.00) | 0.01<br>(0.01;<br>0.01) | 0.01<br>(0.01;<br>0.02) |

### Mean phenotypic plasticity in yolk components along the laying sequence: population level

5α-dihydrotestosterone, all three antioxidants (vitamin E and both carotenoids), the proportion of ω-6 PUFA and the ratio of total ω-6/total ω-3 PUFA decreased over the laying sequence (Table 1; Figure 2). The other steroid hormones (androstenedione, testosterone, and corticosterone) and the proportion of ω-3 PUFA did not change, while the proportion of saturated fatty acids increased from the first to the last egg. There was moderate support for the proportion of monounsaturated fatty acids to increase over the laying sequence (posterior probability = 0.97).

**Figure 2.**
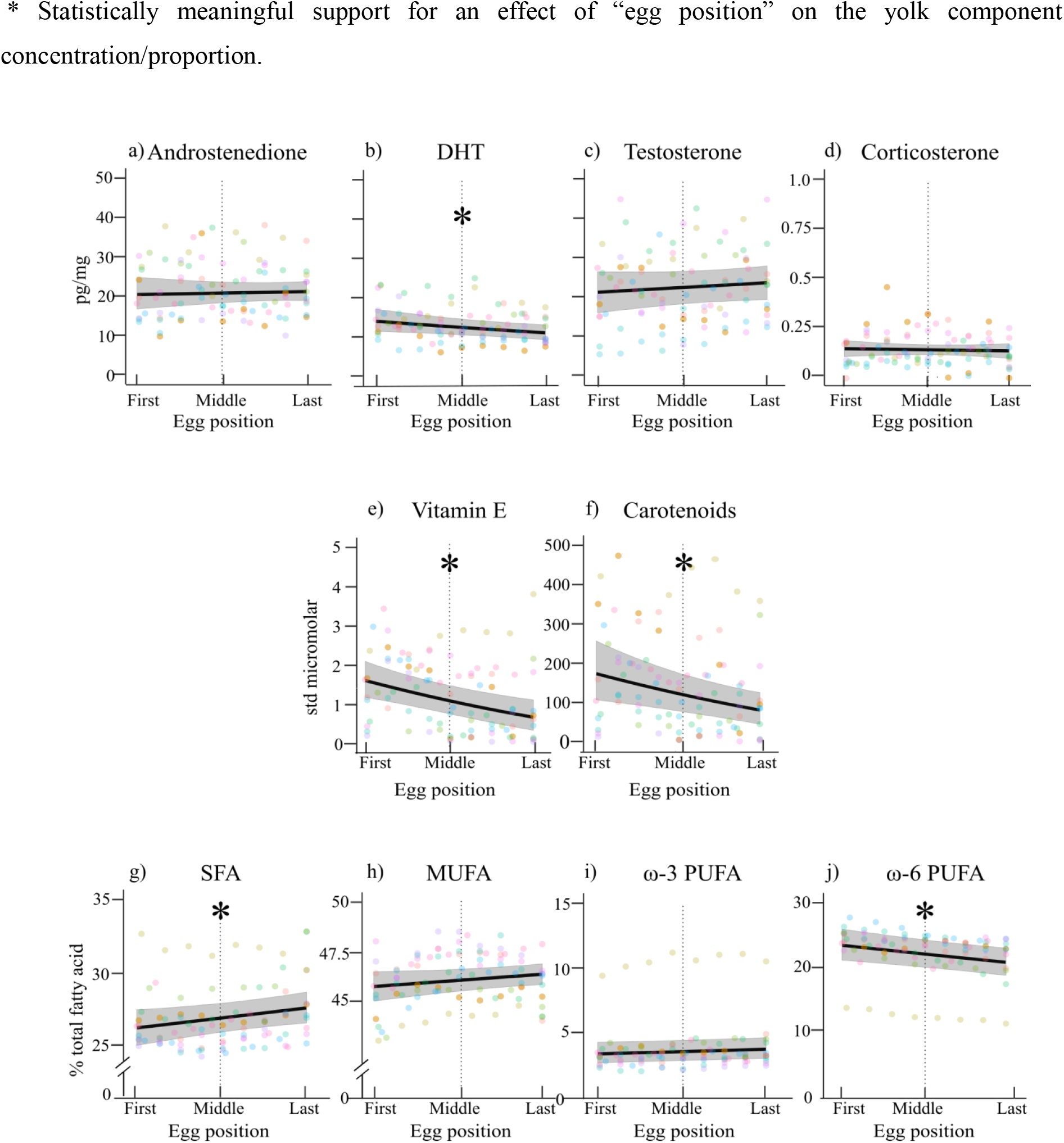
Mean phenotypic plasticity of steroid hormones (a-d), antioxidants (e-f), and fatty acids (g-j) along the laying sequence in great tit egg yolks. Egg position is provided as the egg number relative to total clutch size for each nest. Note that in all statistical analyses the laying sequence was included as a continuous variable and references in x-axis to egg position within the laying sequence are only for illustrative purposes. Filled circles show raw data for each egg; each colour indicates a different nest. The black solid line represents average concentrations or proportions of components, with 95% credible intervals indicated in grey shading. (b) DHT, 5α-dihydrotestosterone; (e) Vitamin E, αtocopherol; (f) Carotenoids, sum of lutein and zeaxanthin; (g) SFA, saturated fatty acids; (h) MUFA, monounsaturated fatty acids; (i - j) PUFA, polyunsaturated fatty acids.

### Variation in yolk components along the laying sequence: female level

Females not only differed in mean concentrations/proportions of yolk components allocated, but also in their phenotypic plasticity over the laying sequence (variation in elevation and slope, respectively; Table 1, Figure 3). Furthermore, the mean trait value and the slope were negatively correlated (i.e., showed a “fanning-in” pattern) in five of the 11 components measured: androstenedione, testosterone, both carotenoids (i.e., lutein and zeaxanthin) and monounsaturated fatty acids (Figure 3, Table 1). Females that on average allocated low levels of androstenedione, testosterone and monounsaturated fatty acids were more plastic, i.e., increased the concentrations/proportions of these yolk components along the laying sequence more strongly than did females with high average concentrations/proportions. In contrast, the decrease in carotenoid concentrations along the laying sequence was more pronounced in females that allocated higher average concentrations of carotenoids into their yolks. For vitamin E, the correlation between elevation and slope was positive (Table 1), but this correlation was driven by one female (Figure 3c; uppermost line) and therefore this result should be treated with caution. The elevation and slope of allocation for the rest of the components were only weakly correlated (Table 1).

**Figure 3.**
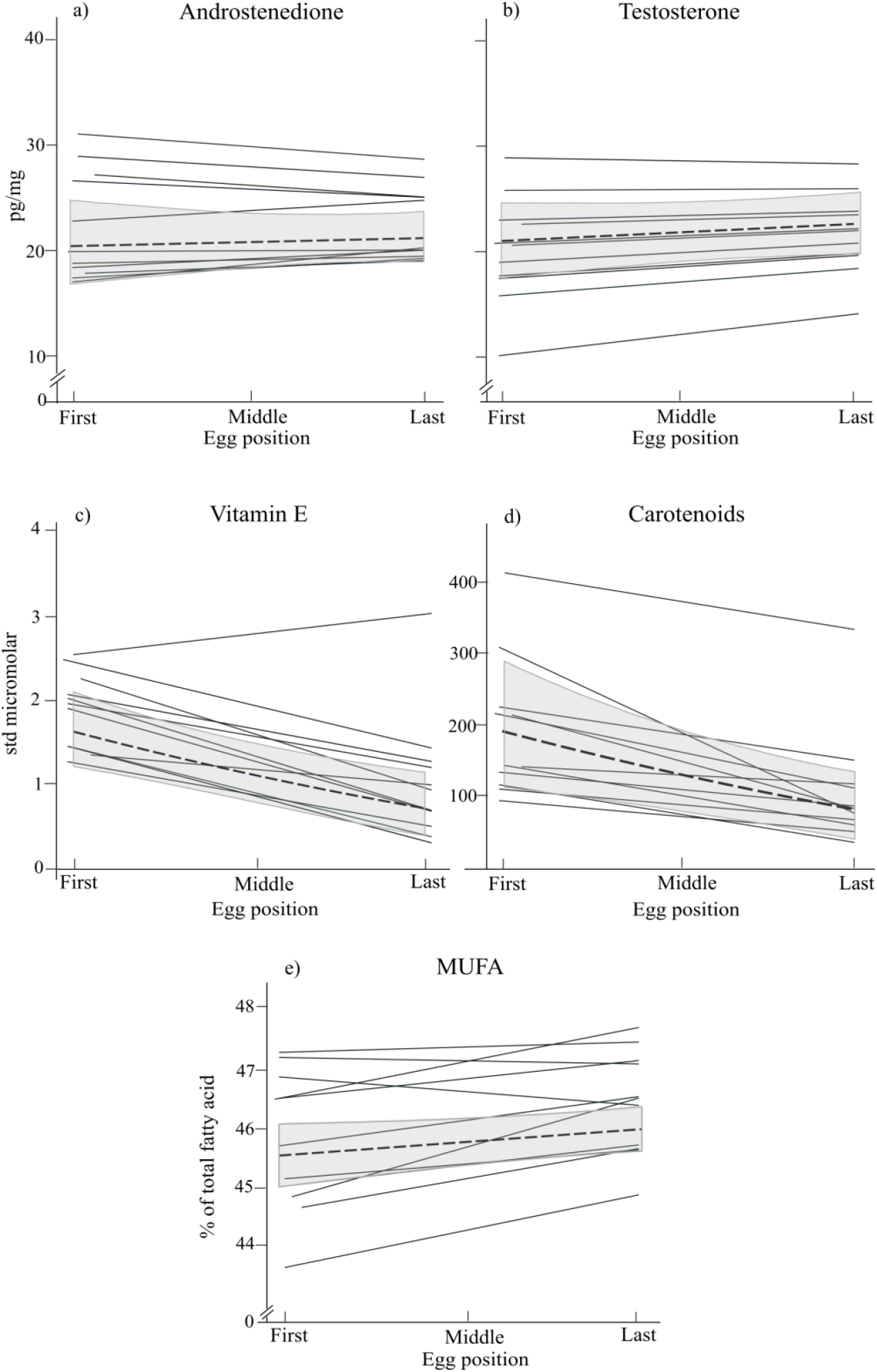
Reaction-norm plots of steroid hormones (a-b), antioxidants (c-d), and fatty acids (e), illustrating among-female variation in yolk allocation patterns along the laying sequence. Plots are shown only for those yolk components where elevation and slope of allocation n were correlated. Egg position is shown as the egg number relative to total clutch size for each nest. Note that in all statistical analyses the laying sequence was included as a continuous variable and references in x-axis to egg position within the laying sequence are only for illustrative purposes. Gray lines indicate different nests. Mean population concentrations or proportions of components (dashed black line) and 95% credible intervals (gray shading) are also shown as a reference. (c) Vitamin E, α-tocopherol; (d) Carotenoids, sum of lutein and zeaxanthin; (e) MUFA, monounsaturated fatty acids.

For the vast majority of the yolk components (10 of 11), female adjusted repeatabilities were higher than 0.30 (Figure 4). The fatty acid groups had the highest repeatability values (ω-3 PUFA: *R* = 0.90; ω-6 PUFAs: *R* = 0.92), followed by the antioxidants, which all had values of repeatability approaching *R* ~ 0.40. Steroid hormones showed the widest range of repeatabilities, with 5α-dihydrotestosterone exhibiting a high (*R* = 0.64), testosterone and androstenedione a moderate (*R* = 0.43 and *R* = 0.33 respectively), and corticosterone a relatively low repeatability (*R* = 0.18).

**Figure 4.**
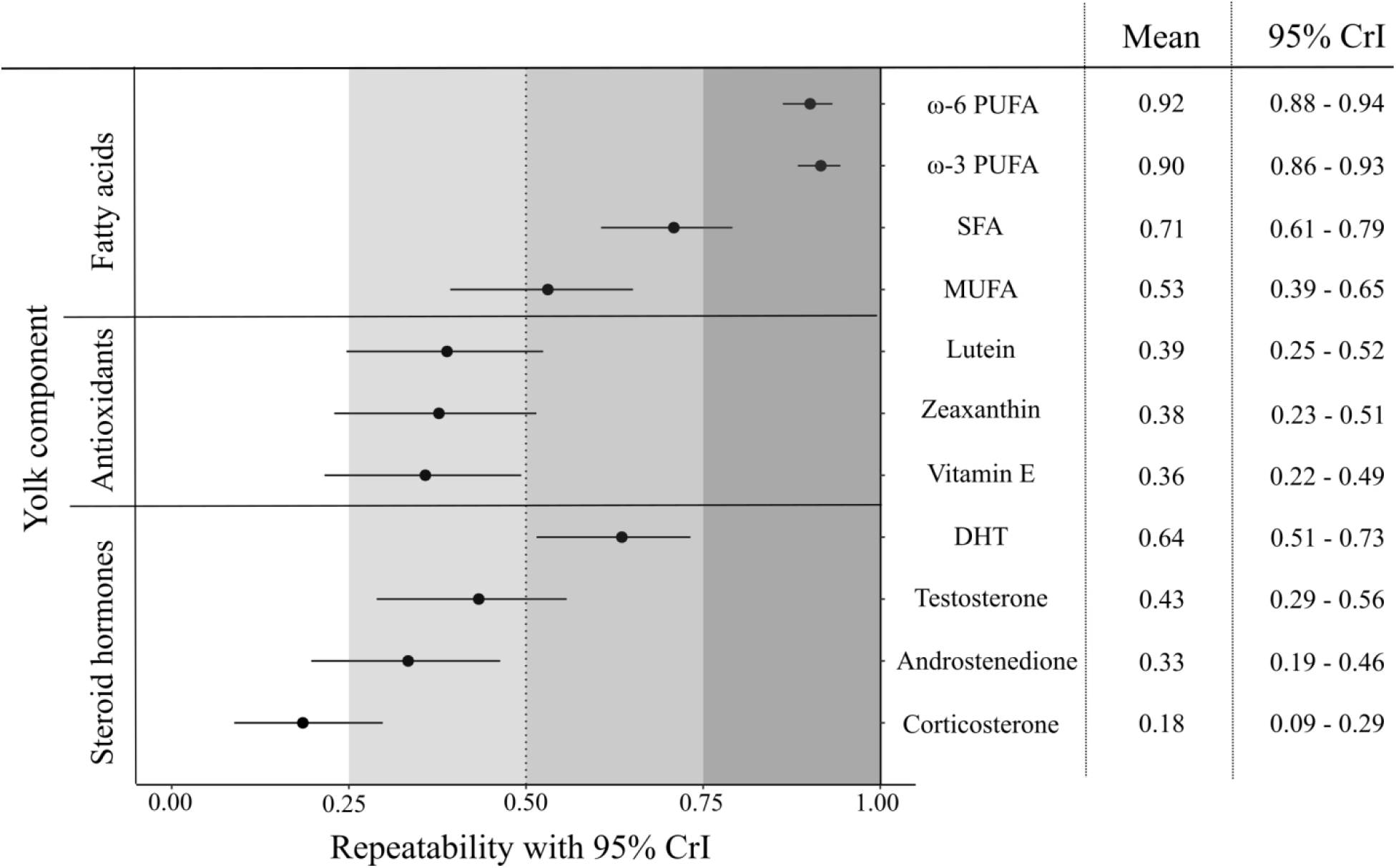
Adjusted repeatabilities of fatty acids (top), antioxidants (middle), and steroid hormones (bottom) of yolk components among females. Repeatability estimates (black circles in the graph and values in the left column of the table) and 95% credible intervals (CrI, horizontal lines in graph and numbers in right column of table) were obtained from linear mixed-effects models. Gray shading of increasing intensity indicates increases in repeatability. DHT, 5α-dihydrotestosterone; Vitamin E, αtocopherol; SFA, saturated fatty acids; MUFA, monounsaturated fatty acids; PUFA polyunsaturated fatty acids.

## Discussion

We quantified 11 yolk components and analyzed their variation along the laying sequence by acknowledging the multi-level nature of female resource allocation. At a population level, the concentrations/proportions of five resources in the yolk varied along the laying sequence: first-laid eggs generally contained higher concentrations of 5α-dihydrotestosterone, antioxidants and proportions of ω-6 PUFA, and lower proportions of saturated fatty acids than last-laid eggs (Table 1; Figure 2). This result implies that in general the physiological environment is rather different for offspring developing in first *versus* last eggs. Individual females allocated yolk components over the course of laying in a pattern that was not necessarily the same as observed at the population level (Table 1; Figure 3). Females differed in their mean allocation of components (i.e., differences in the elevation of reaction norms) indicating that some females allocated on average higher amounts of components into their eggs than other females, while they also varied in their plastic response over the laying sequence (i.e., differences in the slope of reaction norms; Table 1; Figure 3). In addition, for some yolk components these two parameters were correlated. Finally, egg component allocation was repeatable, i.e., the concentration/proportion of most yolk components was more similar among eggs from the same female than among eggs from different females (Figure 4). Overall, these results show that at both population- and female-level, the physiological environment of the offspring will be different depending on the egg from which they develop. However, even if females are plastic in their allocation of components along the laying sequence, those eggs laid by the same mother are more similar to each other as compared to the eggs laid by a different mother.

### Mean phenotypic plasticity in yolk components along the laying sequence: population level

For yolk hormones, consistent species-specific patterns of allocation along the laying sequence have been described for avian eggs (reviewed by Groothuis et al. 2005, Gil 2008). However, opposing patterns for the same hormone have also been described for different populations of the same species (reviewed by Groothuis et al. 2005, Gil 2008). For example, in free-living great tits an increase in androstenedione concentrations from first to last egg has been reported (Tschirren et al., 2004; Lessells et al., 2016), whereas in a study on great tits from selection lines the mean phenotypic plasticity showed opposing patterns depending on behavioral traits (Groothuis et al. 2008), and in our study androstenedione did not change. Further, a decrease in 5α-dihydrotestosterone along the laying sequence was reported by Lessells et al. (2016) and our study, but not in the other two above-mentioned studies. Testosterone concentrations increased over the laying sequence in Tschirren et al. (2004) and Groothuis et al. (2008; only females from the ‘bold’ line), but no such trend was observed by Lessells et al. (2016) and the current study. Finally, while in a previous study yolk corticosterone increased (Lessells et al. 2016), we found no change along the laying sequence. This lack of agreement among different great tit populations could be explained, on the one hand, by female quality (e.g., body condition; Supplementary materials Appendix 1, Table A4), environmental conditions (e.g., mean ambient temperature; Table 1), consistent individual differences between females (“personality”; Ruuskanen et al. 2018), and social factors, such as territory quality and male condition or personality (Remeš 2011, Ruuskanen et al. 2018), which all may influence female yolk allocation. In addition, opposing population trends could also be explained by different patterns of individual female plasticity (see below).

In the current study, all three antioxidants (i.e., vitamin E, lutein, and zeaxanthin) decreased along the laying sequence, confirming previous studies showing that last-laid eggs generally have lower concentrations of antioxidants in birds (Royle et al. 1999, 2003, Hõrak et al. 2002, Blount et al. 2002, Rubolini et al. 2011; but see Török et al. 2007). Animals cannot synthesize vitamin E and carotenoids *de novo*, therefore these antioxidants have to be obtained from the diet (Surai and Speake 2008) and can then be allocated into the egg yolk (from where the developing embryo will absorb them; reviewed by Yigit et al. 2014). These antioxidants may represent a limiting resource for the mother (Møller et al. 2000), and indeed, females in better body condition on average allocated higher concentrations of vitamin E into their eggs (Supplementary materials Appendix 1, Table A4). However, lesser black-backed gull (*Larus fuscus*) females supplemented with a diet rich in carotenoids had higher carotenoid concentrations in plasma and yolk, but they also decreased yolk carotenoid concentrations over the laying sequence in a similar way as non-supplemented birds (Blount et al. 2002). Since the last-laid eggs in our study were not inferior to first-laid eggs in terms of egg mass (which increased over the laying sequence; Supplementary materials Appendix 1, Table A1), these findings suggest that the observed decline in antioxidants cannot be attributed solely to female depletion in nutrients and other resources over the laying period.

Eggs were collected at a time when great tits were changing their diet from mainly feeding on seeds to predominantly feeding on invertebrates (caterpillars) which, compared to seeds, are particularly rich in saturated fatty acids, the ω-3 PUFA α-linolenic acid, and carotenoids (Isaksson and Andersson 2007, Andersson et al. 2015, Isaksson et al. 2015). In addition, a strong correlation between fatty acid levels of ingested food and fatty acids in yolk has previously been established (Lin et al. 1991, Hulbert and Abbott 2011, Twining et al. 2016). Thus, the changes in fatty acid composition along the laying sequence reported here (i.e., an increase and decrease in the proportion of saturated fatty acids and ω-6 PUFA, respectively) may be explained by an increase in caterpillar availability over the course of the breeding season, and a lower reliance on seeds, which are richer in ω-6 PUFAs. In line with this idea, lay date had a statistically positive effect on saturated fatty acids and ω-3 PUFA proportions (Table 1). However, other factors may also contribute to the observed patters given that the laying period in great tits can be quite short (mean ± SD: 8.45 ± 1.13 days in this study). For instance, in contrast to PUFAs, saturated and monounsaturated fatty acids can be biosynthesized *de novo* by animals, and fatty acids can also be selectively mobilized from internal stores to plasma (Raclot 2003, Price et al. 2008), suggesting that female condition at the start of laying may also play a role for the fatty acid allocation. Lastly, to date only three studies have documented mean phenotypic plasticity in fatty acid proportions along the laying sequence in free-living birds (Bourgault et al. 2007; Toledo et al. 2016). In contrast to our findings, a recent study on several populations of great tits in the UK found no variation in fatty acid composition along the laying sequence (Toledo et al. 2016). In that study, however, fatty acid composition was analyzed only for the 2^nd^ to the 5^th^ eggs in the laying sequence, which is equivalent to analyzing yolks only from the first to the middle eggs in our study. After re-analyzing our data with only first to middle eggs (n = 49 eggs), the fatty acid proportion still changed along the laying sequence (results not shown), thus indicating that it is unlikely that the difference in the position of the eggs analyzed explains the differences in the patterns of allocation obtained in the two studies. On the other hand, Toledo et al. (2016) collected one egg per nest. Our finding that yolk fatty acid composition in general is more similar in eggs from the same mother compared to those from other mothers (i.e., high repeatability; see below) could explain the differences in allocation patterns reported in these two studies.

Finally, it is important to bear in mind that ethical considerations limited our sample size (i.e., of entire clutches collected), potentially reducing our statistical power to identify the environmental or internal factors underlying population-level variation in yolk steroid hormones, antioxidants and fatty acids allocation.

### Co-secretion of yolk components

To date, most studies investigating the fitness consequences of yolk allocation patterns focused on single groups of yolk components. This approach has been important for understanding the ways in which females can generate transgenerational phenotypic plasticity in the offspring by allocating certain substances into their eggs, as well as the evolutionary consequences. However, such studies may misinterpret the fitness benefits of single yolk components because we now know that different classes of components are allocated at the same time, often targeting the same phenotypic traits in the offspring (e.g., Treidel et al. 2013). We simultaneously quantified yolk steroid hormones, antioxidants and fatty acids, components that are known to affect growth, immune responses and oxidative stress of offspring, but sometimes in opposite directions. Although our study cannot directly address patterns of co-secretion of certain components because of limitations in sample size (n=11 clutches), we have observed some tantalizing patterns of co-occurrence in our study population that merit further investigation. For instance, first-laid eggs were high in 5α-dihydrotestosterone concentrations and ω-6 PUFA proportions. Both of these components are essential to promote embryo development, but they can also potentially increase the concentration of reactive oxygen species and thereby induce oxidative stress (Pamplona et al. 2002, Larsson et al. 2004, Alonso-Alvarez et al. 2007, Hulbert and Abbott 2011). However, first-laid eggs also had high antioxidant concentrations, which can buffer oxidative stress (e.g., Royle et al. 2001, Surai et al. 2001, Watson et al. 2018). These findings raise the question of whether selection promotes females to allocate eggs with a particular yolk composition, for example by co-secreting substances that have growth-enhancing but oxidative-stress inducing effects (like androgens and PUFAs) together with substances that can mitigate oxidative damage like antioxidants. Supporting this idea, a positive correlation between testosterone and vitamin E concentrations in the yolk of 75 bird species was recently reported (Giraudeau and Ducatez 2016). Detailed studies of the co-secretion of several yolk components are now required to address the question of whether natural selection operates on female allocation patterns to balance the costs and benefits of allocating substances to the offspring. Furthermore, in addition to quantifying the fitness costs and benefits incurred by females during the egg-laying stage (i.e., through resource acquisition, synthesis, allocation, etc.), future studies should also determine the costs and benefits that arise later at the offspring stage – to the mother (e.g., by having to provision more demanding offspring), to the offspring (e.g., by having a suboptimal phenotype given the environmental and social circumstances), and to the female’s partner (e.g., by having to increase investment into parental care).

### Variation in yolk components along the laying sequence: female level

Our findings provide the first evidence that females consistently differ in average amounts of yolk components *and* in their plasticity of allocation along the laying sequence (i.e., in their slope). This result could explain the existence of divergent mean phenotypic plasticity (i.e., at a population level) found in studies on different populations of the same species (reviewed by Groothuis et al. 2005, Gil 2008). Our current understanding of the mechanisms behind female variation in allocation patterns is limited. Female mean yolk androgen deposition shows moderate heritability (e.g., great tits, Ruuskanen et al. 2016). However, whether mean deposition of antioxidants and fatty acids is also partially explained by heritable variation still remains unknown. Furthermore, which genetic factors may underlie female phenotypic plasticity also represents an important unanswered question. Ecological parameters that affect female physiological condition like prevailing climatic conditions, female quality or population density could also potentially alter female yolk allocation along the laying sequence. Genetic and non-genetic sources can simultaneously affect both components of the reaction norm (i.e., the variation in the average amount of yolk components and female plasticity), thus contributing to the overall among-female variation observed.

The elevation-slope coefficients that we obtained in our analyses for yolk components like androstenedione, testosterone, carotenoids, vitamin E and MUFAs should be interpreted with caution because of the low sample sizes (Martin et al. 2011, van de Pol 2012). Nevertheless, our analyses allowed us to quantify the covariation between two sources of variation, i.e. the extent and the direction to which the elevation and slope in the allocation of one yolk component were correlated in females (Table 1; Figure 3). For example, a negative correlation between elevation and slope indicates that females that overall allocated a higher concentration of a component also decreased this component’s concentration more strongly along the laying sequence (i.e., were more plastic). Such a pattern could suggest that females experienced a constraint along the laying sequence. Furthermore, the presence of correlations between elevation and slope in yolk components indicates that consequences of such maternal effects cannot be fully evaluated by studying these two sources of variation independently. For those components where female mean allocation and individual plasticity are correlated, fitness consequences (e.g., number of chicks that hatched or fledged) that would be attributed to one source of variation, for example to mean yolk concentrations in testosterone through an analysis of only the elevation of the allocation reaction norm, might in fact be caused by the other component, i.e., the change in yolk testosterone concentrations along the laying sequence. Lastly, this finding also raises the question of whether females with different reaction norms experience divergent fitness consequences. In other words, do females that on average deposit a higher proportion of e.g., monounsaturated fatty acids but are less plastic along the laying sequence have higher reproductive success than females that deposit on average lower proportions of that components but are more plastic (Figure 3e)?

Female-level variation in yolk allocation as well as the basis behind such plasticity still is a largely unexplored field of research. Does selection shape the allocation of average levels of egg components, the degree of plasticity over the laying sequence, or the correlation between these two traits? Addressing these questions will require large sample sizes, which might be a limiting factor in this field of research for ethical reasons. However, combining data from different populations and research groups could be a rewarding avenue to overcome this obstacle and increase our knowledge of the evolutionary and ecological forces driving phenotypic female variation in yolk allocation.

### Repeatability in yolk allocation

The repeatability estimates in the present study varied depending on type of yolk resources allocated (Figure 4). Ours is the first study to report (adjusted) repeatabilities for antioxidants and fatty acids in egg yolks. The medium-high repeatabilities observed for these two groups (ranging from 0.36 to 0.92, Figure 4) are perhaps not surprising since great tits usually lay one egg each day, and may have experienced homogeneous environmental conditions within this short time frame. In contrast, the lower repeatability reported here for yolk hormones might be due to the fact that steroid hormones are synthetized by the mother herself (Groothuis and Schwabl 2008, Gil 2008). Within the group of steroid hormones measured, corticosterone concentrations had the lowest repeatability estimates. Plasma corticosterone levels are known to fluctuate over short time scales, and its concentration in yolk may be influenced by maternal circulating concentrations (Saino et al. 2005, Groothuis and Schwabl 2008, Pitk et al. 2012). Therefore, the low repeatability estimates obtained for corticosterone might result from variations in maternal plasma concentrations along the laying sequence. Interestingly, while in our study 5α-dihydrotestosterone was the steroid hormone with the highest repeatability estimate (*R* = 0.64), in another recent study on great tits this hormone showed the lowest repeatability value (*R* = 0.29; Lessells et al. 2016). However, care should be exercised when comparing different studies because repeatability is a coefficient between variance explained by female identity in relation to the total variance (Nakagawa and Schielzeth 2010). Differences among females in their ability to synthesize and/or allocate each yolk component, the methodology used to measure each component, or environmental conditions that might increase the residual (unmeasured) variance, can all modify repeatability estimates even when the among-female variance remains the same. We therefore propose that future studies should report both among-female variance and repeatability estimates to allow for a better comparison of female consistency in yolk allocation across populations.

Repeatability estimates are a useful tool for evolutionary ecologists because they enable the quantification of the upper limit to heritability (Boake 1989). These estimates therefore provide information about the potential genetic contribution to the measured phenotype as well as clues as to whether some traits might evolve in response to selection. In our study, repeatability estimates for antioxidants and fatty acids were high. However, this does not necessarily indicate a high heritability in the allocation of these components. High repeatability estimates in our study could also be explained by repeated measurements taken at very short intervals (Araya-Ajoy et al. 2015, Holtmann et al. 2017) and/or by the fact that we adjusted for environmental factors such as lay date and mean ambient temperature. Irrespective of differences in absolute estimates of repeatability, the finding that repeatabilities for almost all yolk components were moderate to high (*R* ≥ 0.30, with the exception of corticosterone, Figure 4) indicates that the developmental environment for offspring of the same mother is largely similar. Importantly, the high repeatability estimates obtained also confirm that the method of analyzing a single egg (ideally the middle egg) from a nest is a suitable way to estimate clutch-level yolk composition in studies of wild populations (e.g., Giordano et al. 2014), a technique that allows to assess the consequences for offspring phenotypes and fitness.

## Conclusions

The present study emphasizes the need to account for female variation in yolk allocation along the laying sequence at multiple levels as a way to increase our understanding on the evolutionary processes that shape female yolk allocation. At a population level, our study shows that the developmental environment provided by mothers is different for offspring developing in first *versus* last eggs for almost half of the 11 components measured. Although not analyzed quantitatively, our study raises the question whether the patterns of allocation observed for steroid hormones, antioxidants and fatty acids are the result of selection favouring a complementary allocation of yolk components. Interestingly, the patterns of allocation at an individual level differed from the general pattern observed at a population level. At a female level, individuals varied among each other in the average allocation of yolk components, in their plasticity along the laying sequence as well as in the correlation between both parameters. In addition, females were remarkably consistent in the allocation of the majority of yolk components, confirming that the method of collecting a single egg from a nest is a suitable way to estimate clutch-level yolk composition in studies of wild populations – at least in those that aim to quantify the consequences for offspring phenotypes. Future studies can now build on these findings and test these patterns and their consequences in other species. It would also be important to analyze whether individual females are consistent in their allocation of yolk components across clutches laid in the same or in different years and if female plasticity along the laying sequence varies across homogeneous *vs*. heterogeneous environments. Since female allocation may be key to understand patterns at a population level, using mixed models to study female allocation at multiple levels opens up promising fields of research.

## Declarations

### Acknowledgements

We are grateful to Sabine Jörg and Nicolás Adreani for their invaluable help in the field, Anna Johansson and Hong-Lei Wang for important contributions to fatty acid extraction and analysis, and Amparo Herrera-Dueñas and Jürgen Kuhn for antioxidant extractions and quantification. We also thank Fränzi Korner and María Moirón for their help and insightful discussions regarding the statistical analysis. We thank Glenn Cockburn and María Moirón for their constructive criticism of previous versions of the manuscript.

## Authors’ contributions

LM conceived the study, conducted the field work, analyzed the data and drafted the manuscript. LM and MH designed the study with input from WG and CI. LM and MT conducted egg and steroid analysis. CI and MNA supervised the antioxidant and fatty acid extractions and analyses. MH, WG, CI and MNA contributed to manuscript preparation. All authors approved of the final version of the manuscript.

## Permits

All experimental procedures were conducted according to the legal requirements in Germany and were approved by the governmental authorities of Oberbayern, Germany (license number 55.2-1-54-2532-25-2015).

